## Supplementary information for "Complex interactions in the life cycle of a simple parasite shape the evolution of virulence"

by Luis M. Silva and Jacob C. Koella

**Table S1.** Longevity, fecundity and virulence evolution.
**Table S2.** Host developmental traits.
**Table S3.** Infection dynamics.
**Table S4.** Virulence metrics and decomposition.
**Figure S1.** Host developmental traits per replicate.

**Table S1. Longevity, fecundity, and virulence evolution.** Longevity was analyzed using a cox proportional hazard with the different regimes as explanatory variables and the replicate as a random factor. Once the survival curves were generated, the maximum hazards for each of the survival curves were extracted using the "muhaz" package [1]. Maximum hazard served as our proxy for virulence, which was then compared across treatments using a linear model with the natural log of the maximum hazard as a dependent variable, while the regime was considered an explanatory factor and the replicate as a random factor. The infection cost in fecundity was calculated only for females who laid eggs (and therefore mated and had access to blood). We calculated the percentual infection cost by subtracting the mean uninfected fecundity from each infected individual fecundity for day 10 and day 15 of adulthood and then dividing the resulting value by the infected individual fecundity and multiplying it by 100. Then, differences due to regime and days post-adult emergence were assessed using a linear model with regime and day post-adult emergence as explanatory variables, as well as their interaction. Moreover, we also included the replicate as a random factor. Further details can be found in the "Statistical analyses" section of the Materials and Methods.

| **Tested effect** | | | |
|---|---|---|---|
| **Model 1a** - Host survival | **$\chi 2$** | ***df*** | ***p*** |
| Regime | 138.82 | 2 | **< 0.001** |
| **Model 1b -** Virulence | **$\chi 2$** | ***df*** | ***p*** |
| Regime | 13.239 | 1 | **< 0.001** |
| **Model 1c - Cost of infection in fecundity** | ***df*** | **F** | ***p*** |
| Regime | 2 | 5.914 | **0.003** |
| Day | 1 | 20.279 | **<0.001** |
| Replicate | 4 | 0.695 | 0.596 |
| Regime : Day | 2 | 1.583 | 0.207 |

**Table S2. Host developmental traits.** Larvae and pupae mortality were assessed as the proportion of dead individuals for each stage. A generalized linear model with a binomial error structure and the replicate as a random factor was used to determine the effect of the regime on each of the developmental stage mortality. Individuals pupated from day 6 to 11 post eclosion, but approximately 77%, 79%, and 82% of them pupated on day eight or nine, for Early, Late, and Stock, respectively. Therefore, we split individuals into two categories, the ones that pupated until day eight and the ones that pupated from day nine onwards, and used a generalized linear model with binomial distribution to measure differences in early pupation. As before, the regime was considered an explanatory variable and the replicate as a random factor. Differences in body size (using wing length as a proxy) were quantified using two linear models: i) the first model tested for differences between infection by selected parasites (i.e., Early and Late) or unselected (i.e., Stock) using selection/no-selection as the explanatory variable and replicate as a random factor; ii) then, we tested for differences due to infection between selected parasites, meaning Early and Late regimes, using a similar model, with the parasite treatment as an explanatory factor and replicate as a random factor. Further details can be found in the "statistical analyses" section of the materials and methods.

| **Tested effect** | **$\chi 2$** | ***df*** | ***p*** |
|---|---|---|---|
| **Model 2a -** Larvae mortality | | | |
| **Selected *vs*. Stock:** | | | |
| Selection | 3.564 | 1 | 0.059 |
| **Early *vs.* Late:** | | | |
| Regime | 3.437 | 1 | 0.064 |
| **Model 2b-** Pupae mortality | | | |
| **Selected *vs*. Stock:** | | | |
| Selection | 2.324 | 1 | 0.127 |
| **Early *vs.* Late:** | | | |
| Regime | 7.890 | 1 | **0.005** |
| **Model 2c -** Early pupation | | | |
| **Selected *vs*. Stock:** | | | |
| Selection | 0.010 | 1 | 0.920 |
| **Early *vs.* Late:** | | | |
| Regime | 14.267 | 1 | **< 0.001** |
| **Model 2d -** Body size | | | |
| **Selected *vs*. Stock:** | | | |
| Selection | 0.027 | 1 | 0.869 |
| **Early *vs.* Late:** | | | |
| Regime | 0.054 | 1 | 0.816 |

**Table S3. Infection dynamics.** The spore production rate was defined as the proportion of females with detectable spores per regime for the first ten days of adulthood. Mosquitoes with detectable spores were then used to estimate spore dynamics from day one to 18 of adulthood. The spore rate was analyzed using a generalized linear model with a binomial distribution. In contrast, we used a linear model for spore dynamics to assess differences in the log-transformed number of spores. In both models, we had regime and day and their interaction as explanatory factors and the replicate as a random factor. Further details can be found in the "statistical analyses" section of the materials and methods.

| **Tested effect** | | | |
|---|---|---|---|
| **Model 3a -** Spore production rate | **χ2** | ***df*** | ***p*** |
| Regime | 4.921 | 2 | 0.085 |
| Day | 114.375 | 1 | **< 0.001** |
| Regime : Day | 24.854 | 2 | **< 0.001** |
| **Model 3b -** Infection dynamics | **χ2** | ***df*** | ***p*** |
| Regime | 41.857 | 2 | **<0.001** |
| Day | 158.934 | 1 | **< 0.001** |
| Regime : Day | 10.993 | 2 | **0.004** |

**Table S4. Virulence metrics and decomposition**. We tested for differences due to parasite regime in virulence, using several host fitness metrics (Model 4a-d). For each of them, we run a linear model with the parasite regime as a fixed factor and replicate it as a random factor. In the presence of a significant effect, we run a posthoc to identify pairwise differences between parasite regimes. After that, we used the maximum hazard from the longevity curves as the main virulence measure and (since the average maximum hazard day was 20) the spore density on days 15 and 18. Spore density was calculated as the individual spore load at days 15 and 18, adjusted by individual wing length. Virulence decomposition analysis was done as in [2]. In brief, exploitation was measured as the natural log-transformed mean spore density from days 15 and 18 for a given regime and replicate, while per parasite pathogenicity was given as the slope of the relationship between natural log-transformed maximum hazard and (i.e., virulence) and exploitation. Lastly, post hoc multiple comparisons were performed with "emmeans", using the default Tukey adjustment across the different models. Infectivity is previously defined as the *number of parasites released from the first host to the next stage, which can be the outside environment or a new host,* according to Silva *et al.* (2025) [3]. Hence, the chronic parasite load of each individual was considered a proxy of infectiousness as it reflects the amount of parasite that will be transferred to the environment in case of host death (particularly in water). The relationship between infectiousness and virulence was then analyzed using a linear model with infectiousness (or mean chronic load at days 15 and 18 of adulthood) as the dependent variable and the log-transformed maximum hazard of every parasite line as the independent variable. Further details can be found in the "statistical analyses" section of the materials and methods.

| **Tested effect** | | | |
|---|---|---|---|
| **Model 4a -** Virulence using larval mortality | ***df*** | **F** | ***p*** |
| Regime | 2 | 13.161 | **0.001** |
| Replicate | 1 | 1.388 | 0.264 |
| Multiple comparisons for Model 4a | ***df*** | ***t*-ratio** | ***p*** |
| Early - Late | 11 | -3.486 | **0.013** |
| Early - Stock | 11 | -1.517 | 0.321 |
| Late - Stock | 11 | 5.003 | **0.001** |
| **Model 4b -** Virulence using pupae mortality | ***df*** | **F** | ***p*** |
| Regime | 2 | 3.310 | 0.075 |
| Replicate | 1 | 1.426 | 0.258 |
| **Model 4c -** Virulence using adult mortality | ***df*** | **F** | ***p*** |
| Regime | 2 | 8.800 | **0.005** |
| Replicate | 1 | 3.940 | 0.073 |
| Multiple comparisons for Model 4c | ***df*** | ***t*-ratio** | ***p*** |
| Early - Late | 11 | -3.383 | **0.016** |
| Early - Stock | 11 | 0.457 | 0.893 |
| Late - Stock | 11 | 3.840 | **0.007** |
| **Model 4d -** Virulence using clutch fecundity | ***df*** | **F** | ***p*** |
| Regime | 2 | 7.669 | **0.008** |
| Replicate | 1 | 5.348 | 0.041 |
| Multiple comparisons for Model 4d | ***df*** | ***t*-ratio** | ***p*** |
| Early - Late | 11 | 2.127 | 0.129 |
| Early - Stock | 11 | -1.784 | 0.220 |
| Late - Stock | 11 | -3.911 | **0.006** |
| **Model 4e -** Exploitation | ***df*** | **F** | ***p*** |
| Regime | 2 | 55.466 | **< 0.001** |
| Multiple comparisons for Model 4e | ***df*** | ***t*-ratio** | ***p*** |
| Early - Late | 12 | -6.086 | **0.001** |
| Early - Stock | 12 | 4.401 | **0.002** |
| Late - Stock | 12 | 10.487 | **<0.001** |
| **Model 4f -** Per parasite pathogenicity | ***df*** | **F** | ***p*** |
| Regime | 1 | 0.441 | 0.531 |
| Day | 1 | 2.866 | 0.141 |
| Regime : Day | 1 | 0.300 | 0.606 |
| **Model 4g -** Infectivity-virulence | ***df*** | **F** | ***p*** |
| ln maximum hazard | 1 | 9.244 | **0.009** |

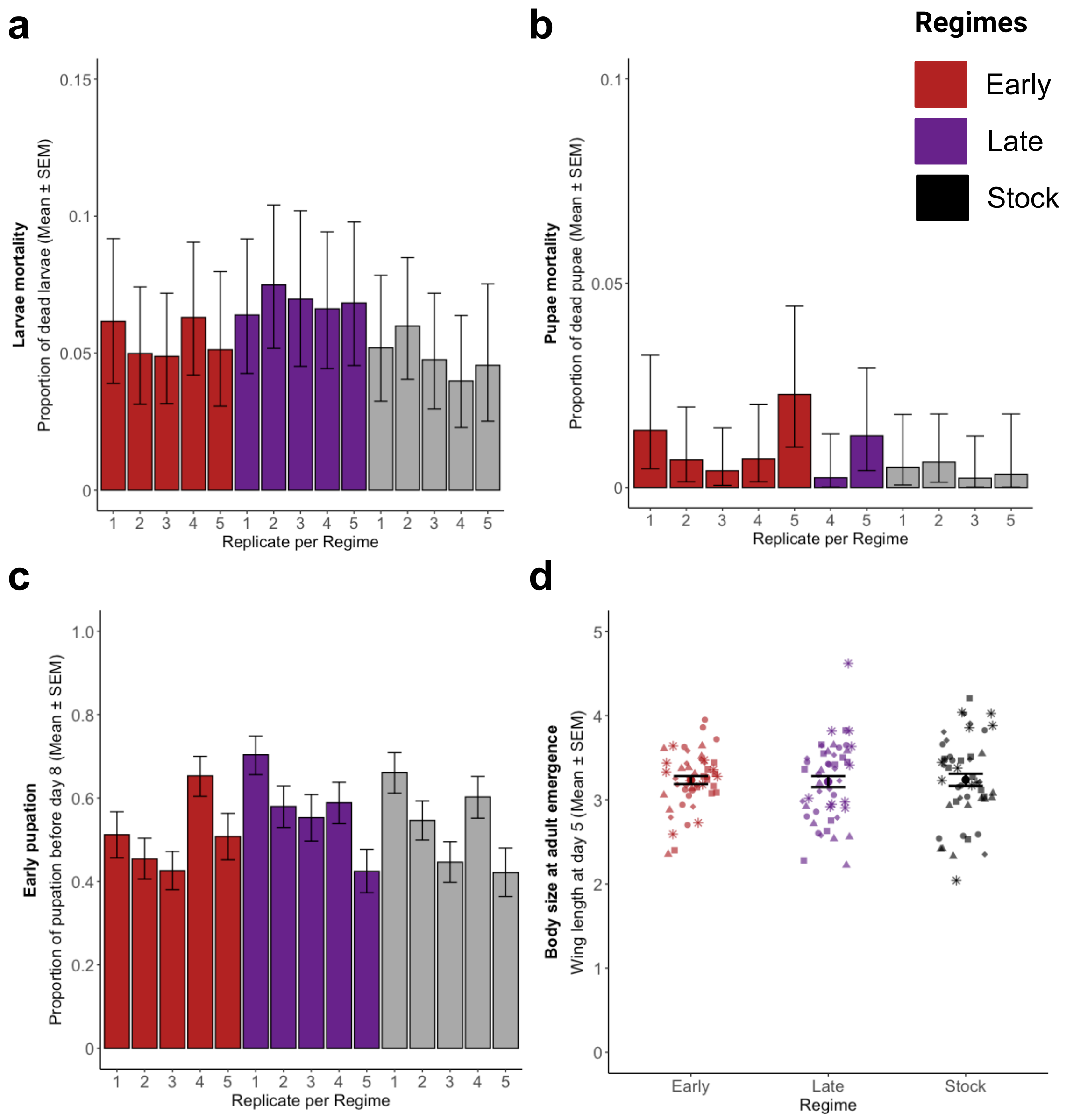


**Figure S1. Host developmental traits per replicate. (a)** Larvae mortality across regimes and replicates. Sample size as follows: Early with 357, 441, 491, 428 and 351: Late with 422, 427, 344, 423 and 395; Stock with 404, 484, 441, 401 and 307 for replicate 1 to 5 respectively. **(b)** Pupae mortality for each regime and replicate. Missing bars represent replicates with no pupae mortality. Sample size was: Early with 335, 419, 467, 401 and 333: Late with 395, 395, 320, 395 and 368; Stock with 383, 455, 420, 385 and 293 for replicate 1 to 5 respectively. **(c)** Early pupation, as the proportion of individuals that pupated until day eight post eclosion. Sample size was the following: Early with 330, 416, 465, 398 and 325: Late with 395, 395, 320, 394 and 363; Stock with 381, 452, 419, 385 and 292 for replicate 1 to 5 respectively. **(d)** Body size at adult emergence using wing length of five-day-old females as a proxy. Each replicate consisted of 10 individuals, illustrated by a different dot shape.
